# Enhanced respiratory electron dissipation by immunometabolites promotes mycobacterial biofilm longevity

**DOI:** 10.1101/2025.10.08.681301

**Authors:** Kaushik Poddar, Snehal V. Khairnar, Stuti Srivastav, Amitesh Anand

## Abstract

*Mycobacterium tuberculosis* organizes as multicellular structures like granuloma and biofilm within human lungs and these lifestyles are critical for the pathophysiological outcomes. Granuloma has been the hallmark of *M. tuberculosis* infection and has been studied extensively. However, the metabolic interplay dictating biofilm development remains poorly understood. We show that supplementation of immunometabolites, nitrate or fumarate, extends mycobacterial biofilm lifespan dramatically. This longevity is enabled by suppression of dormancy regulons and maintenance of active metabolic state. These results establish that access to alternative electron acceptors directly influences mycobacterial biofilm fate. By linking dormancy suppression to prolonged structural integrity, our study identifies respiratory flexibility as a determinant of mycobacterial biofilm persistence. These findings reveal a central metabolic lever that dictates biofilm survival and open new avenues for targeting mycobacterial biofilms in clinical settings.

## Introduction

*Mycobacterium tuberculosis*, the etiological agent of tuberculosis (TB), persists within host-derived multicellular structures known as granulomas ^1^. In addition to these host-mediated assemblies, mycobacteria can generate self-produced multicellular formations, including cords and biofilms ^2,3^. These diverse structural lifestyles are intricately linked to TB pathophysiology. While granuloma formation remains a defining hallmark of TB, cord formation represents a pivotal event in early pathogenesis, and biofilm development during infection has been shown to enhance mycobacterial virulence ^2,4,5^. Notably, such multicellular structures exhibit elevated drug tolerance and are implicated in the emergence of antibiotic resistance.

Biofilms are a widespread mode of bacterial community organization that follow an aging cycle ^6^. Their development begins when cells attach to a suitable surface and proliferate to form a mature structure, in which they become embedded within an extracellular matrix. As biofilms mature, they acquire a multilayered architecture, with bacterial populations adopting metabolic states tailored to the local physico-chemical environment. Despite mounting evidence for the clinical importance of the mycobacterial biofilm lifestyle, its mechanisms of development and dispersal remain poorly understood.

The metabolic interplay between the host and microbes play a crucial role in influencing colonization outcomes. Pathogens can exploit host microenvironments and metabolites to enhance their survival ^7^. During infection, host-derived nitric oxide is converted to nitrate under conditions of high oxidative stress within the inflamed niche. Nitrate has been shown to promote mycobacterial growth in macrophages ^8,9^. Another macrophage derived pro-inflammatory metabolite, fumarate, accumulates significantly in lungs and can impact microbial pathophysiology ^10^. Both these immunometabolites can influence mycobacterial energy metabolism and pathogenic outcomes ^11^. Despite these insights, the effect of such host metabolites on the physiology of mycobacterial biofilms, structures that confer persistence and drug tolerance, remains poorly understood. Defining this relationship is therefore critical to elucidating metabolic adaptations that underlie biofilm-associated tuberculosis.

We examined these immunometabolites for their influence on mycobacterial biofilm survival. In this study, we demonstrate that metabolites reinforcing respiratory electron dissipation fundamentally reshapes biofilm physiology by: (a) nearly doubling biofilm lifespan, (b) suppressing the dormancy regulon to extend longevity, and (c) sustaining metabolic activity to overcome energetic constraints.

## Results and discussion

### Modulation of mycobacterial biofilm by immunometabolites

In Sauton’s medium, *M. smegmatis* establishes a robust biofilm at the air–liquid interface, reaching maturity within five days. Beyond this point, the pellicle gradually loses structural integrity and ultimately sinks to the bottom of the well, a transition that likely reflects the biofilm dispersal (Figure 1A) ^12^.

**Figure 1:**
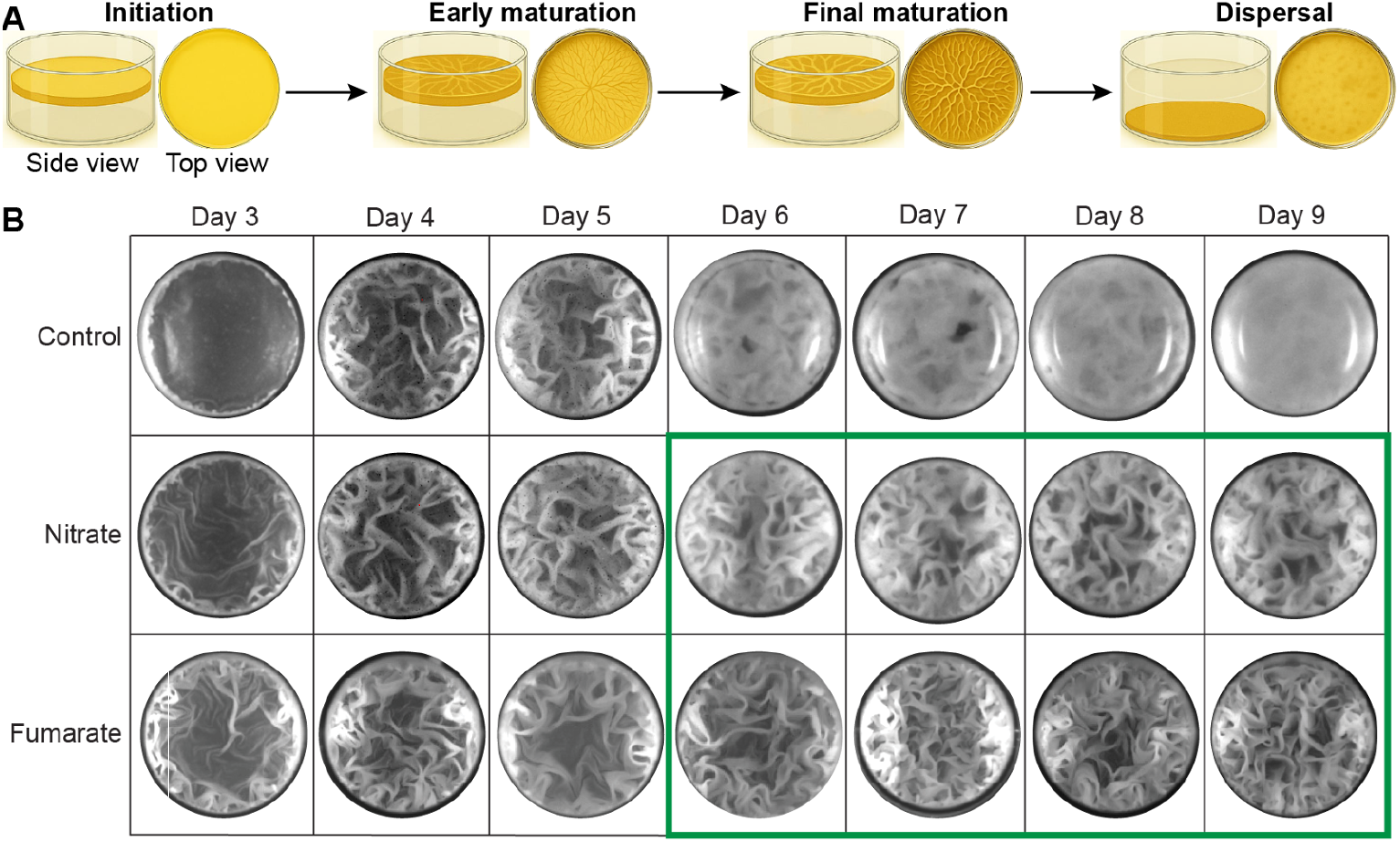
Influence of nitrate and fumarate on the life-span of mycobacterial biofilm. (A) A schematic generated using ChatGPT (OpenAI) showing various stages of in vitro mycobacterial biofilm. (B) Effect of sodium nitrate and sodium fumarate supplementation on *M. smegmatis* biofilm morphology (top view). Columns are for the age of the biofilm in days and rows are for metabolites that are supplemented (1% weight/volume) in the biofilm culture.

We inoculated *M. smegmatis* in biofilm forming media with nitrate and fumarate. We observed that both the supplements hastened the biofilm development (Figure 1B). Interestingly, fumarate and nitrate showed remarkable influence on biofilm lifespan. The untreated biofilms typically disperse by day six, whereas nitrate or fumarate supplementation prolonged biofilm integrity beyond nine days (Figure 1B). The longevity influence was specific to biofilm as we did not observe any significant effect on planktonic cultures (Supp. Figure 1). Mature biofilms can restrict oxygen diffusion across their layers, leading to pronounced hypoxia in the core population ^13,14^. Both fumarate and nitrate can function as alternate electron acceptors and facilitate electron dissipation ^15,16^.

The pronounced extension in biofilm lifespan prompted us to examine whether these metabolites influenced biofilm architecture or cellular survival. We performed further analyses using nitrate which is an established terminal electron acceptor for mycobacteria. Although scanning electron micrographs revealed no major alteration in cellular organization, nitrate-treated biofilms exhibited markedly improved cell viability (Figure 2A, 2B). The mitigation of the decline in live-to-dead cell ratio upon nitrate supplementation suggests that nitrate respiration enhances redox balance and supports hypoxic survival within the biofilm.

**Figure 2:**
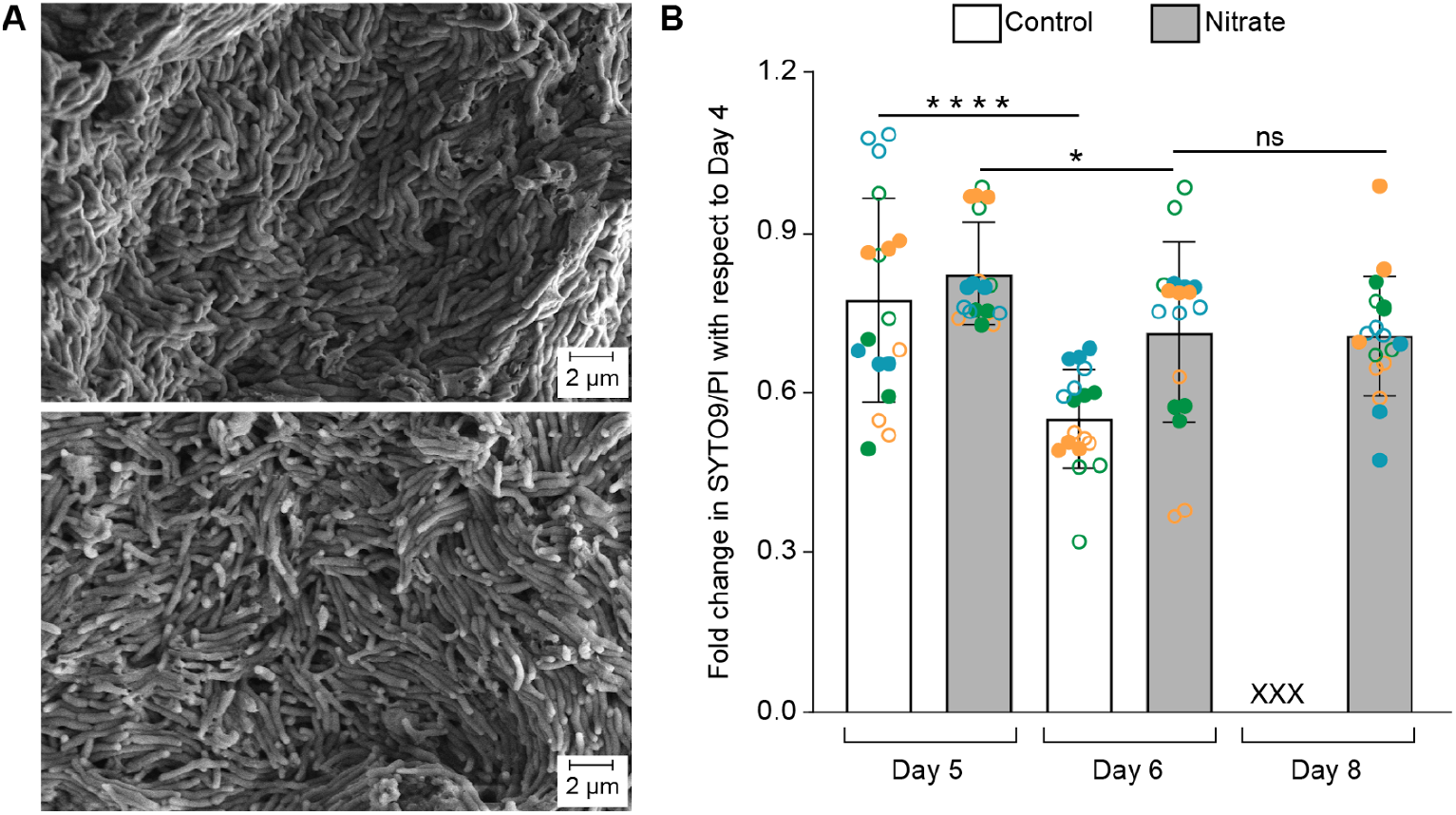
Influence of nitrate on the cells within mycobacterial biofilm. (A) Scanning electron micrograph of *M. smegmatis* biofilms at 10kX magnification (control on the top and 1% sodium nitrate supplemented on the bottom). (B) Fold change in ratio of fluorescence readings of SYTO green-fluorescent nucleic acid stain (SYTO9) and propidium iodide (PI). The unsupplemented biofilm disperses by day 6, so there is no experimental value for day 8. Circles differing either in color or pattern represent different biological replicates. Error bars are showing standard deviation and Unpaired t-test has been used for statistical analysis (ns: non significant, *: p-value < 0.05 and ****: p-value < 0.0001).

### Transcriptional rewiring governing mycobacterial biofilm aging

To investigate the mechanistic link between nitrate supplementation and biofilm aging, we performed temporal profiling of the transcriptome. We observed a clear segregation of age-associated transcriptomes along the first principal component (Figure 3A). Interestingly the transcriptome of nitrate-supplemented biofilm localized closer to the samples from an earlier day. We performed a single-sample gene set enrichment analysis to evaluate pathway activity across samples ^17^. Transcriptomic profiles segregated into five clusters based on the expression of pathway-associated genes (Supp. Figure 2). As expected, cluster 3—the largest cluster, enriched for nitrogen metabolism—showed elevated activity in nitrate-supplemented samples. Interestingly, in the second-largest cluster (cluster 2), the Z-scores of day 5 nitrate-supplemented samples closely resembled those of day 4 controls, indicating a nitrate-mediated delay in biofilm aging.

**Figure 3:**
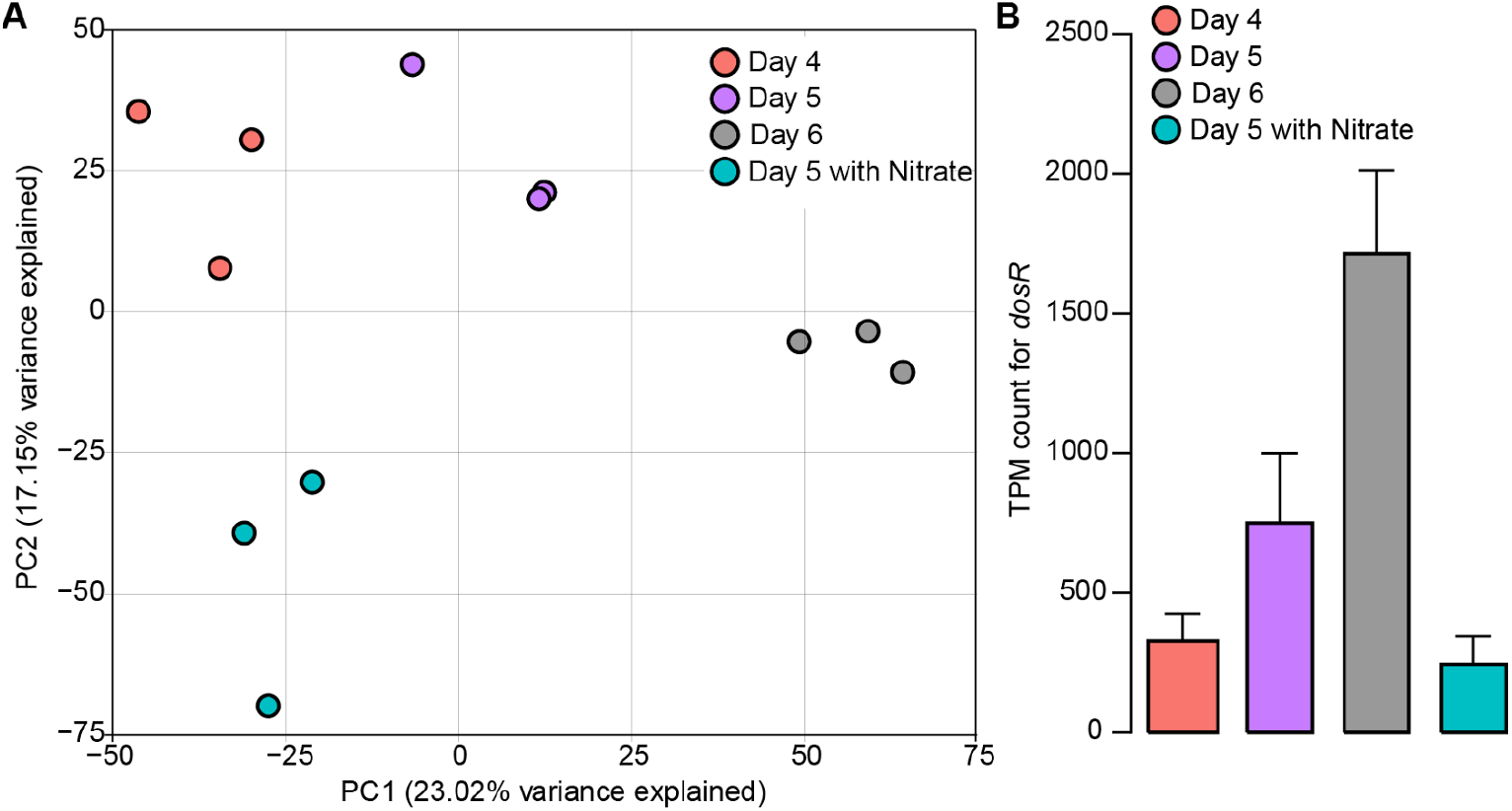
Nitrate induced transcriptional rewiring to prolong *M. smegmatis* biofilm life-span. (A) Transcriptome based distribution of biofilm samples along first and second principal component axes. (B) Temporal expression profile of the dormancy regulator *dosR* in the biofilm (error bar represents standard deviation).

Mycobacterial cells tend to enter a dormant state to survive low oxygen conditions ^18^. The biofilm of *M. tuberculosis* shows decreased expression of dormancy regulon (DosR regulon) ^19^. However, a temporal transcriptomic profiling of aging *M. bovis* BCG biofilm showed progressive increase in the expression of *dosR* ^20^. We observed a transcriptional upregulation in the DosR regulon in *M. smegmatis* biofilm (Figure 3B and Supp. Figure 3). Remarkably, nitrate supplementation suppressed the activation of dormancy in maturing biofilm, thereby, extending the lifespan by providing a conducive metabolic state.

The rewiring in carbon metabolism and respiratory pathways allows environment-specific tailoring of cellular physiology. *M. tuberculosis* shifts to a non-replicating persistent (NRP) state when ambient oxygen concentration decreases gradually ^21^. This drug tolerant state is a hallmark of TB granuloma and results from a decrease in metabolic activities ^22^. The metabolic gene expression pattern of hypoxic NRP cells showed a remarkable overlap with dispersing biofilm cells, with a shift to fermentative energetics (Figure 4A) ^23^. Similarly, there was a temporal increase in the expression of high affinity dioxygen reductase-cytochrome bd oxidase-potentially in an attempt to maintain some respiratory activity under oxygen deprivation (Figure 4B). The reinforcement of electron dissipation through nitrate resulted in lesser reliance on cytochrome bd oxidase and the expression went down to levels as in oxygen -sufficient early biofilm stage.

**Figure 4:**
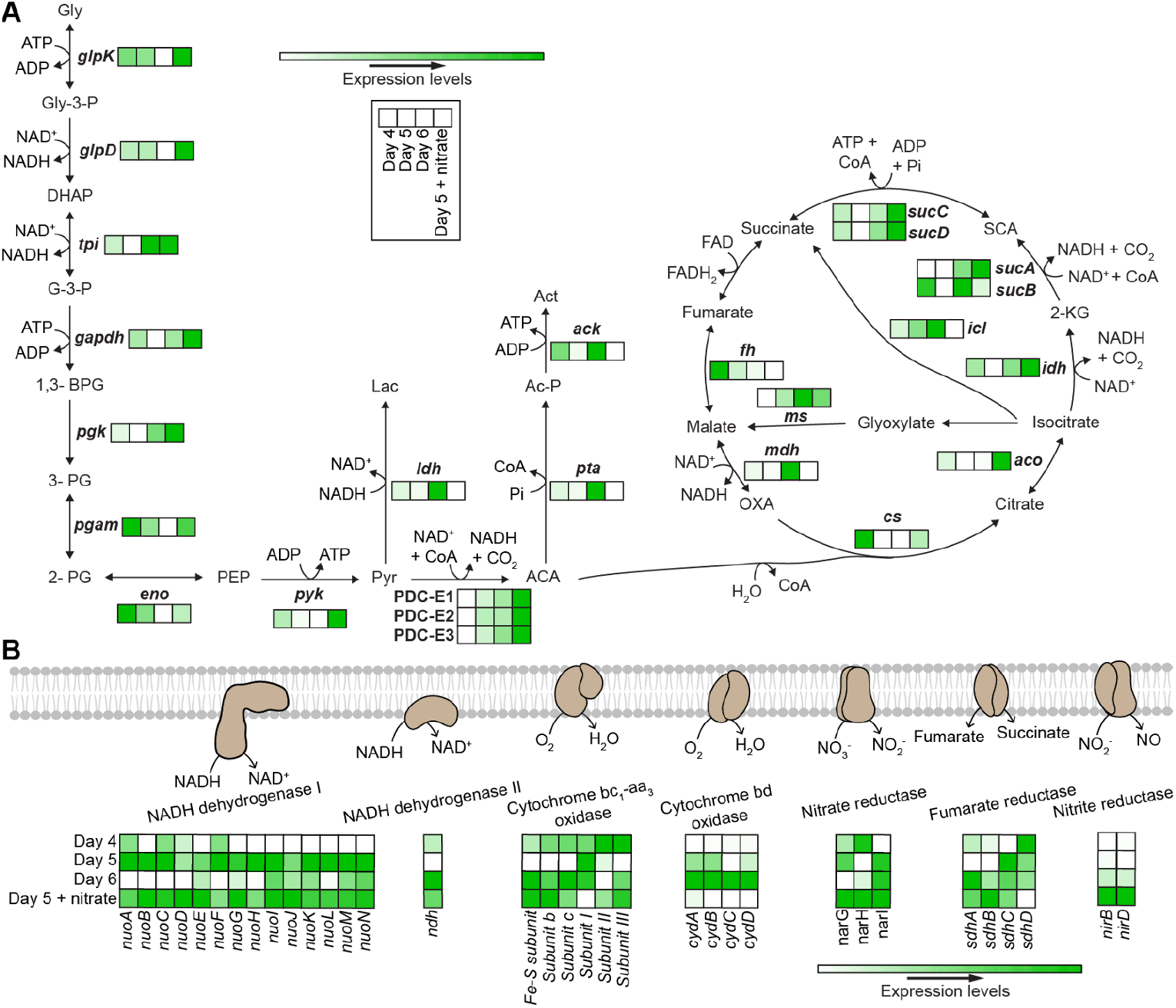
Nitrate induced transcriptional rewiring in metabolic pathways to prolong *M. smegmatis* biofilm life-span. Expression profile of the genes involved in the (A) central carbon metabolism reactions and (B) electron transport system. The expression profile is an average of TPM (transcript per million) counts from three biological replicates. The detailed TPM counts are reported in Supp. Sheet 1.

## Concluding remarks

Our data suggest impaired electron dissipation as a factor causing dispersal of mycobacterial biofilm and mitigating this stress promotes lifespan extension. Given that pathogens encounter a dynamic host milieu enriched with diverse metabolites, shifts in chemical balance—such as the elevated nitrate-to-nitrite ratio favoring *Pseudomonas aeruginosa* colonization in bronchiectasis—underscore the importance of host-derived cues in shaping infection ^24^. *M. tuberculosis* also colonizes lung tissues and forms biofilms, dissecting how microenvironmental components govern its growth and persistence is critical. Notably, modulation of non-replicating persistence signatures may not only alter disease trajectory but also influence therapeutic efficacy.

## Materials and methods

*Mycobacterium smegmatis* mc^2^155 (ATCC-700084) was used in this study. All reagents used, except SYTO green fluorescent nucleic acid stain (Invitrogen), propidium iodide (Invitrogen) and TRIzol (Invitrogen), were from Sigma.

### Planktonic and biofilm growth profiling

Tecan Spark microplate reader was used for growth profiling. The cultures were incubated at 37°C in shaking condition in Sauton’s media containing 0.05% Tween 80. Sodium nitrate and sodium fumarate were supplemented to the media as required. Biofilms were set-up using protocol as described previously ^12^.

### Scanning electron microscopy

The media from biofilm wells were removed carefully. After washing once with 1X PBS, 2.5% glutaraldehyde (made in 1X PBS) was added and incubated overnight at 4°C in the dark for fixation. Glutaraldehyde was then removed and again washed with 1x PBS. Samples were then treated with 1% OsO_4_ for 1 hour. After removing the OsO_4_, samples were washed again with 1X PBS. Finally dehydration of the samples were done by sequential washing with 50%, 70%, 80%, 90% and 100% ethanol. Finally after drying, samples were loaded onto stubs and gold coated for 1 minute at 10mA and imaged using Zeiss Ultra 55 plus.

### SYTO9/PI assay

Biofilm cells were harvested from one well and washed with 1X PBS. 5 uM SYTO9 and PI (made in 1X PBS) were added to the cells and incubated in the dark for 20 minutes. Cells were again washed with 1X PBS to remove any unbound dye. 200 uL of the samples were added to three wells for each sample in black colored, clear bottom 96-well plates. The fluorescence intensity was measured using Tecan Spark multimode microplate reader.

### RNAsequencing

RNA was extracted using Qiagen miRNeasy Mini kit (Cat. No. 1038703) and ribodepletion by NEBNext rRNA depletion kit (Cat. no. NEB E7850X). The NEBNext Ultra II Directional RNA Library prep kit for Illumina (Cat. no. E7760L), was used to construct double-stranded cDNA libraries and sequenced using Illumina NextSeq 2000.

RNA-seq data were processed as described in ^25^, except that reads were mapped to the *M. smegmatis* (NC_008596) reference genome. The batch correction of RNA-seq count data was done using ComBat-seq ^26^.

### Pathway activity analysis by ssGSEA

Single-sample gene set enrichment analysis was used to quantify pathway activity in individual transcriptomic samples ^17^. The analysis was implemented using the GSEApy Python package ^28^ on TPM-normalized, batch-corrected expression data, with gene sets curated from KEGG pathways relevant to *M. smegmatis* ^*29*^. Enrichment scores were standardized using Z-score normalization to facilitate direct comparison across conditions. The mean Z-scores of the three biological replicates, with the standard error of the mean, were then used to perform hierarchical clustering of pathways. This analysis yielded five distinct clusters: Cluster 1 (379 genes), Cluster 2 (458 genes), Cluster 3 (654 genes), Cluster 4 (63 genes), and Cluster 5 (236 genes).

## Supporting information

Supp. Sheet 1

## Acknowledgments

This work was supported by the Ramalingaswami Re-entry Fellowship of the Department of Biotechnology, Government of India (BT/RLF/Re-entry/70/2020) and Tata Institute of Fundamental Research, Department of Atomic Energy, Government of India (19P0120) to Amitesh Anand. We acknowledge the support of the electron microscopy facility of TIFR, Mumbai, India.

## Author contributions

Conceptualization and Supervision: A.A.; Experimentation: K.P. and S.S.; Analysis: K.P., S.K., and A.A.; Manuscript writing: A.A. and K.P.

## Data availability

RNAseq data supporting this study are deposited at GEO (GSE305991).

