## Supplementary figures and tables for "Enhanced respiratory electron dissipation by immunometabolites promotes mycobacterial biofilm longevity"

**Supplementary information**

**Supplementary Figures:**

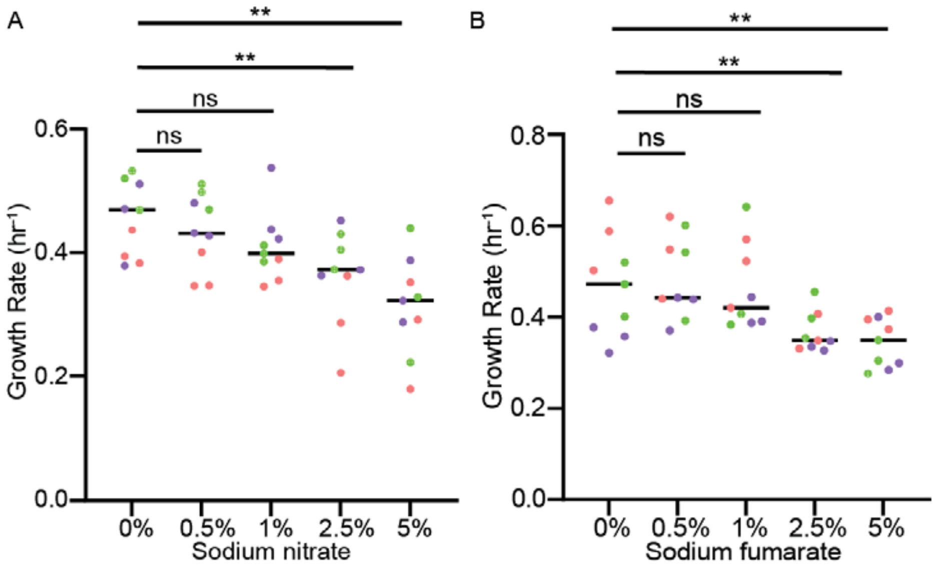


**Supplementary Figure 1:** Planktonic growth profiling of *Mycobacterium smegmatis* with different concentrations of sodium nitrate (A) and sodium fumarate (B). Each color represents three technical replicates of one biological replicate. Mann-Whitney U test was used for statistical analysis (ns: non significant, **: p-value ≤ 0.005).

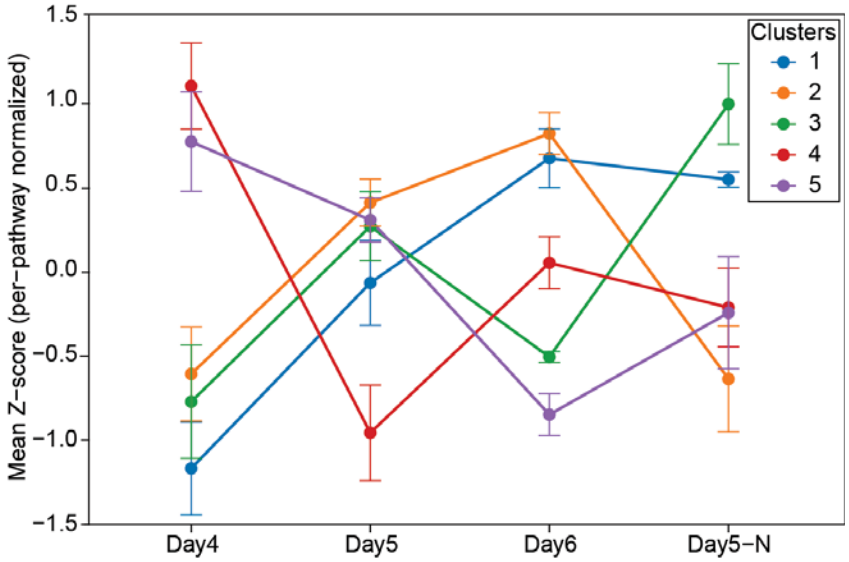


**Supplementary Figure 2:** Pathway activity clusters across biofilm conditions. Mean Z-scored single-sample gene set enrichment analysis (ssGSEA) profiles (cluster centroids) are shown for five pathway clusters across Day 4, Day 5, Day 6, and Day 5 with nitrate. The error bar here represents standard error of the mean. Detailed list of pathways with their genes, ssGSEA scores and Z-score normalized enrichment scores with cluster assignments are given in Supp sheet 1.

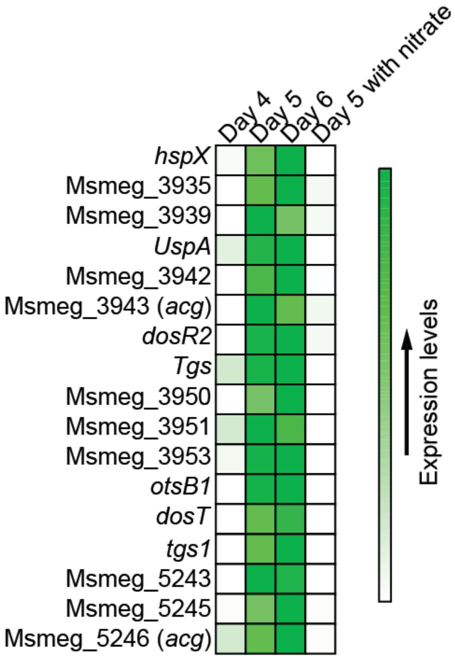


**Supplementary Figure 3:** Expression of the DosR regulon in the biofilm of *M. smegmatis*. The expression profile is an average of TPM (transcript per million) counts from three biological replicates.

**Content of the supplementary sheet:**

List of detailed TPM counts used to generate the expression profiles of DosR regulon, central carbon metabolism and electron transport system genes.

ssGSEA pathway analysis of biofilm samples: KEGG pathways – list of all pathways and their constituent genes used in the ssGSEA analysis. ssGSEA enrichment scores – raw enrichment values computed for each pathway in each sample. Z-score normalized enrichment scores for all pathways (individual replicates and replicate-averaged profiles), with cluster assignments from hierarchical clustering.
